# Distinct roles for the Charcot-Marie-Tooth disease-causing endosomal regulators Mtmr5 and Mtmr13 in axon radial sorting and Schwann cell myelination

**DOI:** 10.1101/843219

**Authors:** Anna E. Mammel, Katherine C. Delgado, Andrea L. Chin, Alec F. Condon, Jo Q. Hill, Sue A. Aicher, Yingming Wang, Lev M. Fedorov, Fred L. Robinson

## Abstract

The form of Charcot-Marie-Tooth type 4B (CMT4B) disease caused by mutations in myotubularin-related 5 (*MTMR5*; also called *SET Binding Factor 1; SBF1*) shows a spectrum of axonal and demyelinating nerve phenotypes. This contrasts with the CMT4B subtypes caused by *MTMR2* or *MTMR13* (*SBF2*) mutations, which are characterized by myelin outfoldings and classic demyelination. Thus, it is unclear whether MTMR5 plays an analogous or distinct role from that of its homolog, MTMR13, in the peripheral nervous system (PNS). MTMR5 and MTMR13 are pseudophosphatases predicted to regulate endosomal trafficking by activating Rab GTPases and binding to the phosphoinositide 3-phosphatase MTMR2. In the mouse PNS, Mtmr2 was required to maintain wild type levels of Mtmr5 and Mtmr13, suggesting that these factors function in discrete protein complexes. Genetic elimination of both Mtmr5 and Mtmr13 in mice led to perinatal lethality, indicating that the two proteins have partially redundant functions during embryogenesis. Loss of Mtmr5 in mice did not cause CMT4B-like myelin outfoldings. However, adult *Mtmr5^-/-^* mouse nerves contained fewer myelinated axons than control nerves, likely as a result of axon radial sorting defects. Mtmr5 levels were highest during axon radial sorting, whereas Mtmr13 levels rose as myelin formed, and remained high through adulthood. Our findings suggest that Mtmr5 and Mtmr13 ensure proper axon radial sorting and Schwann cell myelination, respectively, perhaps through their direct interactions with Mtmr2. This study enhances our understanding of the non-redundant roles of the endosomal regulators MTMR5 and MTMR13 during normal peripheral nerve development and disease.

## INTRODUCTION

During the development of the peripheral nervous system (PNS), Schwann cells associate with bundles of mixed diameter axons. In a process known as radial sorting, Schwann cells segregate large diameter axons (>1 μm) from smaller axons, which remain in bundles (1, 2). Once sorted into 1:1 associations with Schwann cells, large caliber axons are wrapped in a specialized, multilayer membrane sheath called myelin (3). Myelin facilitates rapid, saltatory nerve impulse conduction by clustering sodium channels at nodes of Ranvier, and by reducing axonal capacitance (4). Small caliber axons (<1 μm) remain unmyelinated and are supported by Schwann cells in Remak bundles (2). Both myelinating and Remak Schwann cells provide metabolic support to maintain axon integrity (3, 5–8). Both radial sorting and subsequent myelination require coordinated cytoskeleton reorganization, and receptor signaling from the adaxonal and abaxonal Schwann cell surfaces (1, 3). The precise control of downstream signaling is thought to be mediated in part by receptor trafficking through the endo-lysosomal pathway (9–11).

Genomic studies have strongly suggested a link between abnormal endosomal trafficking and demyelinating Charcot-Marie-Tooth (CMT) disease (12). CMT is the most common inherited neurological disorder, affecting 1 in 2500 people (13). Axonal and demyelinating forms of CMT are defined clinically, and are typically caused by mutations that initially affect axons and Schwann cells, respectively (14). CMT type 4 (CMT4) is characterized by autosomal recessive inheritance and demyelination (15). Nearly half of the human proteins that, when mutated, cause CMT4 are proposed to regulate membrane trafficking through endo-lysosomal compartments. These proteins include dynamin 2 (*DNM2*), factor-induced gene 4 (*FIG4*), FGD1-related F-actin-binding protein (*FRABIN*), lipopolysaccharide-induced tumor necrosis factor-alpha factor (*LITAF*), SH3 domain and tetratricopeptide repeats 2 (*SH3TC2*), myotubularin-related 2 *(MTMR2*)*, MTMR5,* and *MTMR13* (12). Several of these proteins, including FIG4, FRABIN, MTMR2, MTMR5, and MTMR13, control endo-lysosomal trafficking by enzymatically modifying or binding to phosphoinositides (PIs). Despite the generally broad expression of these genes in mammalian tissues, mutations therein primarily impact the PNS, suggesting that myelinating Schwann cells are highly sensitive to disruptions in endo-lysosomal membrane trafficking (12, 16).

CMT4B results from mutations in MTMR2, MTMR5 (also known as SET Binding Factor 1; SBF1), or MTMR13 (SBF2) (Fig. 1) (17–19). Myotubularins are a large family of phosphoinositide 3-phosphatases that regulate membrane trafficking through endosomes and lysosomes (20). This family of proteins includes both catalytically active and inactive phosphatase enzymes. MTMR2 contains an active phosphatase domain, whereas MTMR5 and MTMR13 contain catalytically inactive phosphatase domains (20, 21). MTMR5 and MTMR13 share 59% protein identity and possess the same unique set of protein domains, including a DENN domain that actives Rab GTPases, and a coiled-coil motif that facilitates association with MTMR2 (Fig. 1). MTMR2 and MTMR13 are thought to function as a membrane-associated complex that regulates PI levels and drives Rab GTPase activation to influence the trafficking of key Schwann cell receptors through endosomes (22–25). However, it remains unclear if MTMR5, which also binds avidly to MTMR2 (26), plays an analogous role to MTMR13 in the PNS.

**Figure 1:**
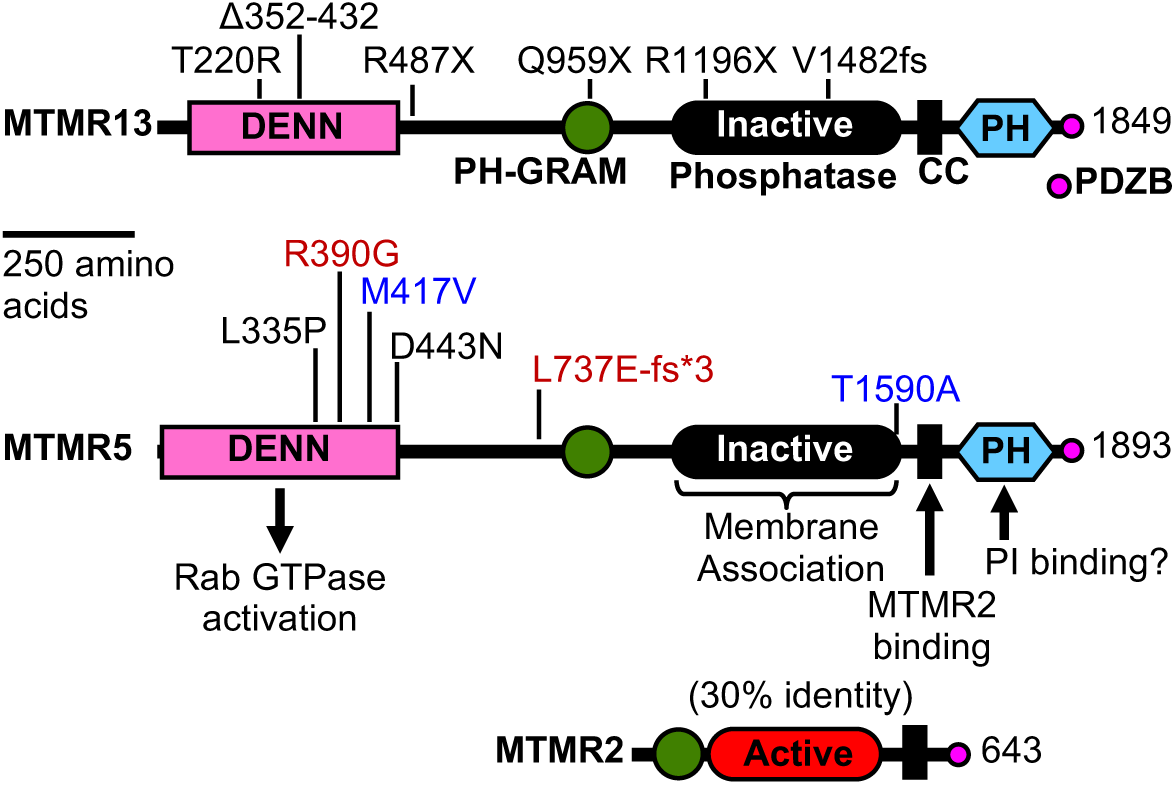
CMT-causing mutations in the MTMR13 and MTMR5 proteins. All mutations are recessive, consistent with a loss-of-function mechanism of pathogenesis. Functions attributed to specific domains of MTMR13/5 are indicated. MTMR5 mutations M417V-T1590V and R390G-L737E-fs*3 are shown in blue and red, respectively, to indicate compound heterozygosity in the corresponding patients. The MTMR13 mutation T220R was identified in a patient with Griscelli syndrome. MTMR5 shares 59% identity with MTMR13. The inactive phosphatase domains of MTMR5 and MTMR13 are about 30% identical to the PI 3-phosphatase domain of MTMR2. *Abbreviations*: PH, pleckstrin homology; DENN, differentially expressed in neoplastic versus normal; GRAM, Glucosyltransferases, Rab Activators, Myotubularins; PDZB, PDZ domain Binding; CC, Coiled-Coil; PI, Phosphoinositide.

The human nerve pathology associated with homozygous mutations in MTMR2 or MTMR13 is well characterized and features distinctive myelin outfoldings thought to arise from excessive longitudinal myelin growth, as well as demyelination, and severe, presumably secondary, axon loss (17, 19). In contrast, MTMR5-associated neuropathies show broader clinical presentations, which include myelin outfoldings, axon loss, thin myelin, and bands of Bungner (18, 27, 28). To date, five families have been identified with CMT-causing mutations in *MTMR5*; a majority of these patients bear missense mutations within or near the DENN-domain (Fig. 1) (18, 27–30). Thus, it remains unclear whether MTMR5 loss-of-function causes myelin outfoldings analogous to those observed in the nerves of CMT4B1 (MTMR2) and CMT4B2 (MTMR13) patients.

To elucidate the role of Mtmr5 in the PNS, we generated an *Mtmr5* knockout mouse model using CRISPR-Cas9 gene editing, and subsequently assessed the pathological consequences of the loss of Mtmr5 on peripheral nerves. We used biochemical approaches to determine how the three CMT-causing Mtmr proteins physically interact, and inferred how these interactions may relate to the observed nerve phenotypes. Our study indicates that the homologous proteins, Mtmr5 and Mtmr13, both require Mtmr2 to maintain wild type levels of protein abundance in the PNS, but each play distinct roles during axon radial sorting and myelination.

## RESULTS

### A novel deletion allele of *Mtmr5*

Initial characterization of an *Mtmr5* deletion allele in mice was described prior to the discovery that mutations in human MTMR5 cause CMT4B3 disease (31). The previously described *Mtmr5* deletion causes male infertility, but whether these mice develop peripheral nerve defects was not examined (31). We attempted to reestablish this mouse line using intracytoplasmic sperm injection, but were unsuccessful. Therefore, an *Mtmr5* deletion allele was generated using CRISPR-Cas9 mutagenesis and subsequently assessed for nerve defects. To target *Mtmr5*, guide RNAs (gRNAs) were designed to target exon 1 and exon 25 (Fig. 2A). Founder mice were screened for mutations in exon 1 and/or large deletions that removed the sequences between exon 1 and exon 25. A founder mouse was determined to be mosaic for two distinct fusions of exon 1 and exon 25 (mutation 1 and 2) (Fig. 2A). Only mutation 2 was passed to progeny mice (Fig. 2B). This novel *Mtmr5* mutation is predicted to yield a frame-shift that leads to a premature stop codon in exon 25; translation would result in a non-functional six amino acid peptide (Fig. 2A; Supp. Fig. S1).

**Figure 2:**
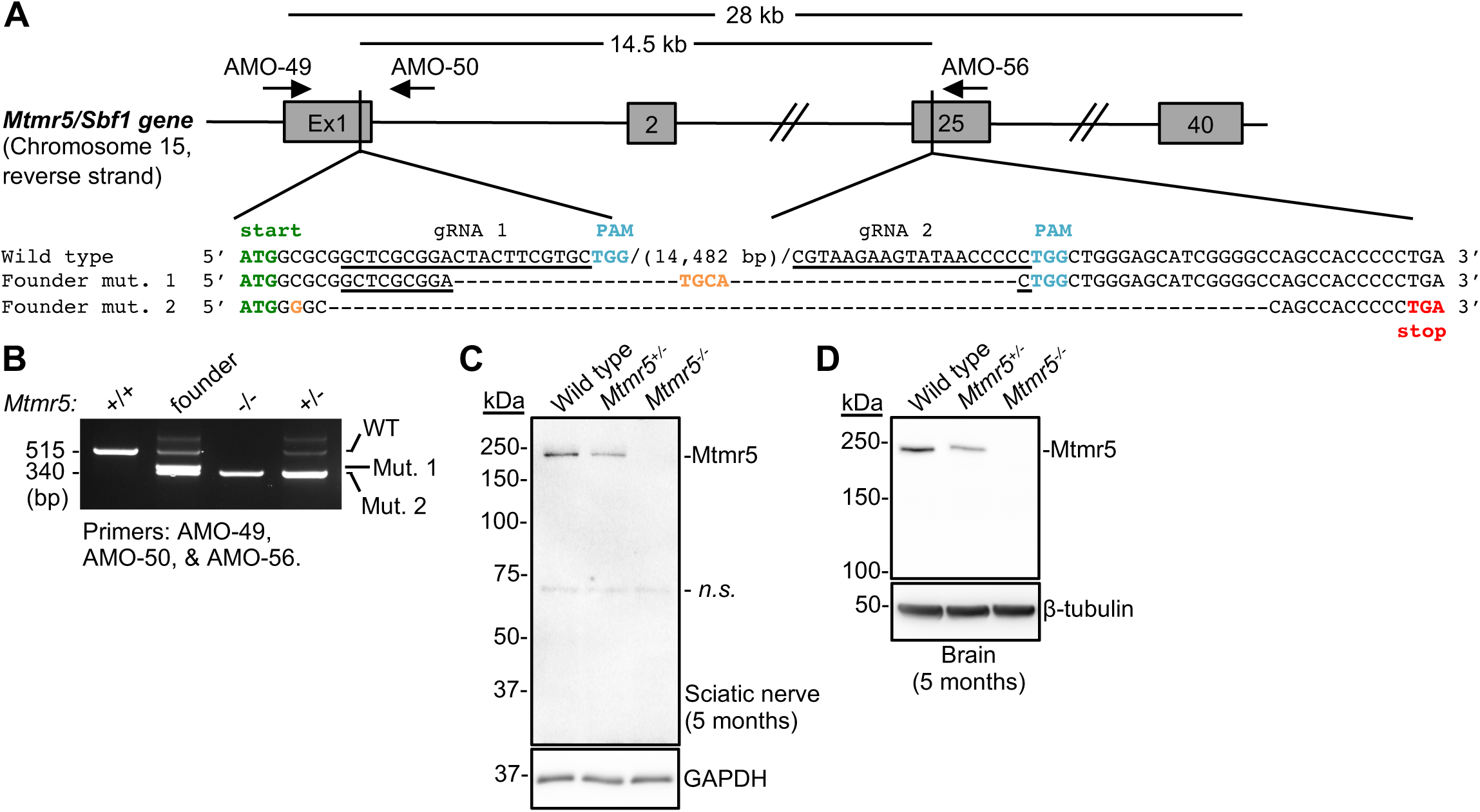
CRISPR/Cas9-mediated disruption of *Mtmr5*. (**A**) A CRISPR-Cas9-mediated 14 kb deletion between exon 1 and exon 25 of *Mtmr5/Sbf1.* The following features are indicated: gRNA target sequences (*underlined*), transcription start site (*green*), PAM sequences (*blue*), premature stop codon (*red*), and additional mutations (*orange*). The CRISPR-Cas9-modified founder mouse was mosaic for two deletions termed mutations 1 and 2 (mut. 1, mut. 2). The genomic locations and orientations of the genotyping primers AMO-49, AMO-50, and AMO-56 are indicated. (**B**) Genotyping of a wild type mouse produces a 515 bp PCR fragment. The F1 generation and all subsequent intercrosses contained mutation 2 deletion exclusively. (**C-D**) Immunoblot analyses of Mtmr5 protein levels from sciatic nerve and brain extracts from 5-month-old wild type, *Mtmr5^+/-^*, and *Mtmr5^-/-^* mice using a C-terminal Mtmr5 antibody. Each sciatic nerve extract was prepared by pooling two nerves. Representative data from one of three independent experiments are shown. The protein loading controls were GAPDH and β-tubulin.

Consistent with the predicted gene disruption, the full-length 208 kDa Mtmr5 protein was undetectable in sciatic nerve and brain exacts from adult *Mtmr5^-/-^* mice (Fig. 2C, D). However, a truncated Mtmr5 protein of 84.8 kDa was detected at a low level in brain extracts from post-natal day 7 (P7) *Mtmr5^-/-^* and *Mtmr5^+/-^* mice (Supp. Fig. S2). We predict that translational initiation at an ATG codon in exon 26 leads to low-level expression of this truncated Mtmr5 protein, at least in neonatal mouse brains. The described, N-terminally truncated Mtmr5 protein was not detected in adult brain or sciatic nerve extracts (Supp. Fig. S2). When expressed, the truncated protein may be non-functional, as it lacks both the DENN and PH-GRAM domains (Fig. 1).

*Mtmr5^-/-^* mice were viable and observed at the frequency predicted by Mendelian genetics. Homozygous mutants of both sexes were about 30% smaller than wild type and *Mtmr5^+/-^* controls (Supp. Fig. S3). This weight difference was observed at weaning and continued throughout the life of the animal. Overt motor or gait deficiencies were not observed in *Mtmr5^-/-^* mice examined at P38 and 17 months (*data not shown*). *Mtmr5^-/-^* males were sterile (Table 1), consistent with previous observations made using a distinct deletion allele (31). The absence of Mtmr5 protein expression, and the sterility of *Mtmr5^-/-^* males strongly suggests that the mutation described here is a *null* or strongly *hypermorphic* allele.

**Table1:** Male sterility in the absence of Mtmr5.

| Breeding Pair |  |  |  |  |
| --- | --- | --- | --- | --- |
| M | x F | Pairs mated (no.) | Litters (no.) | Litter size |
| +/+ | +/- | 3 | 3 | 6 ± 2 |
| +/- | +/- | 9 | 9 | 6 ± 1 |
| +/- | -/- | 3 | 3 | 3 ± 1 |
| -/- | +/+ | 6 | 0 | 0 |

### Biochemical relationships amongst CMT-linked myotubularins

MTMR5 and MTMR13 are both known to bind avidly to MTMR2 in a manner that requires coiled-coil sequences (26, 32). However, it is unclear whether the pseudophosphatases also interact with each other. To test their physical associations, we overexpressed epitope-tagged MTMR5 with MTMR13 and/or MTMR2 in HEK293 cells. MTMR5 and MTMR13 formed a very weak interaction that did not require coiled-coil dimerization, or their mutual binding partner MTMR2 (Fig. 3A-C). The weak interaction between MTMR5 and MTMR13 did not enhance either proteins abundance. In contrast, co-expression of MTMR2 greatly increased the levels of both MTMR5 and MTMR13 (Fig. 3A, B). These data suggest that MTMR5 and MTMR13 may form a very weak association that does not equate to their avid binding of MTMR2.

**Figure 3:**
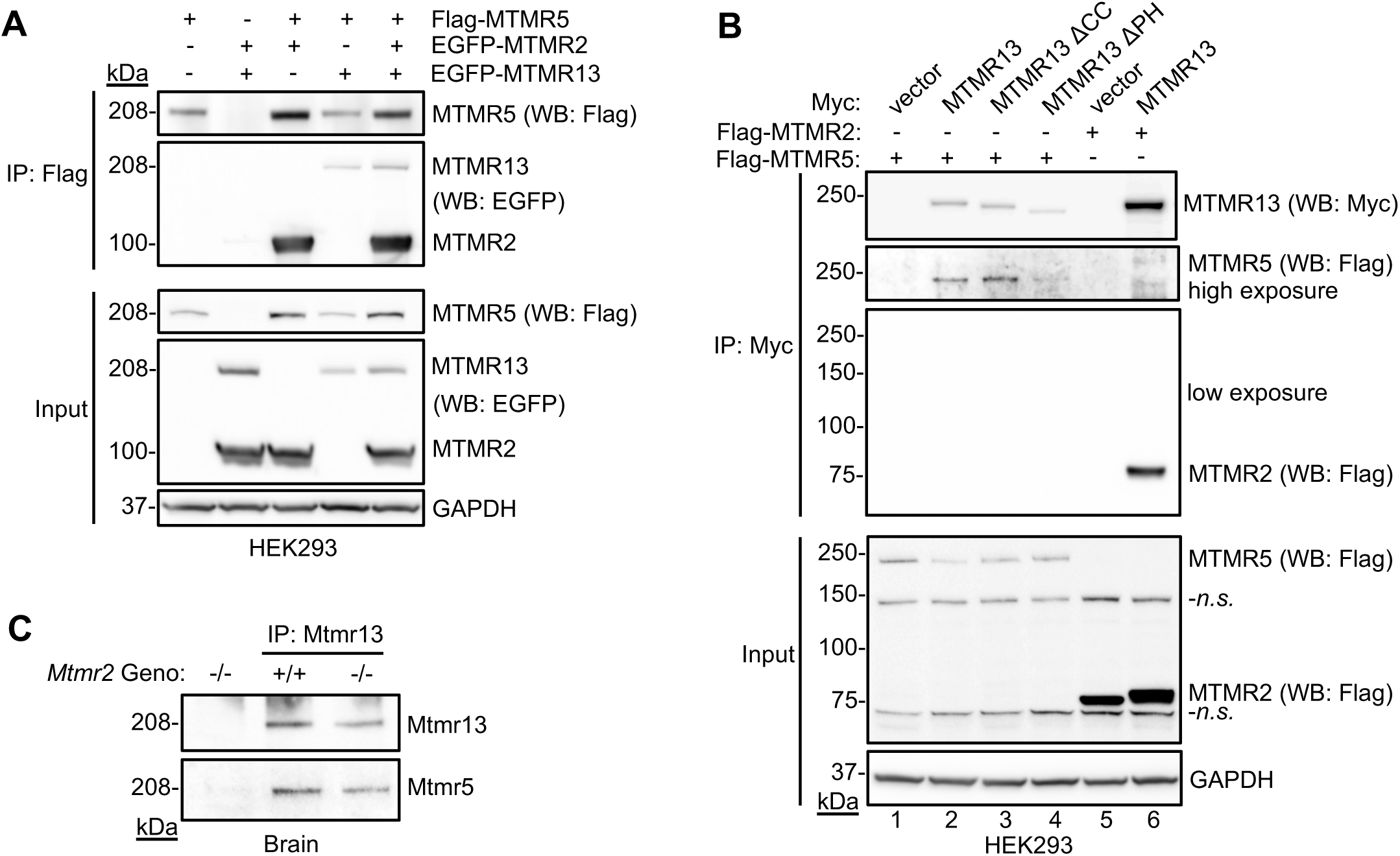
Physical associations between CMT-linked myotubularin proteins. (**A**) HEK293 cells were transfected with vectors expressing the indicated proteins. After 48 h, cell lysates were prepared and analyzed by FLAG immunoprecipitation (IP), SDS-PAGE, and western blotting (WB). Co-expression of EGFP-MTMR2 increased the amount of FLAG-MTMR5 protein in the lysate and IP. FLAG-MTMR5 immunoprecipitated EGFP-MTMR2 much more efficiently than it did EGFP-MTMR13. Representative images from one of two experiments are presented. (**B**) HEK293 cells were transfected with constructs encoding FLAG or c-Myc epitope-tagged versions of the indicated myotubularin proteins. The association between FLAG-MTMR5 and myc-MTMR13 did not require the coiled-coil (CC) motif or the C-terminal pleckstrin homology (PH) domain of MTMR13. Co-expression of FLAG-MTMR2, but not FLAG-MTMR5 increased the recovery of immunoprecipitated myc-MTMR13 (compare lanes 2 and 6). Representative data from one of two independent experiments are shown. Nonspecific (*n.s.*) cross-reactions of the anti-FLAG antibody with proteins present in cell lysates are indicated. (**C**) Coimmunoprecipitation of endogenous Mtmr13 and Mtmr5 from wild type and *Mtmr2^-/-^* mouse brains. The association between Mtmr5 and Mtmr13 was not altered by the absence of Mtmr2. The data from are single experiment are shown.

We previously demonstrated that Mtmr2 and Mtmr13 require each other to maintain wild type protein levels in mouse sciatic nerves (22). To determine if Mtmr2 and/or Mtmr13 proteins were required to regulate Mtmr5 levels, we examined mutant mouse brain and sciatic nerve lysates. Both Mtmr5 and Mtmr13 protein levels were significantly reduced in *Mtmr2^-/-^* nerves (Fig. 4A, B). However, the requirement of Mtmr2 to maintain Mtmr5 at wild type levels was not reciprocal; *Mtmr5^-/-^* sciatic nerve lysates had comparable Mtmr2 levels to wild type nerves (Fig. 4D, E). Mtmr13 loss had no impact on Mtmr5 levels in mouse brain or sciatic nerves, demonstrating that these pseudophosphatases do not require one another to maintain wild type protein levels (Fig. 4A, C).

**Figure 4:**
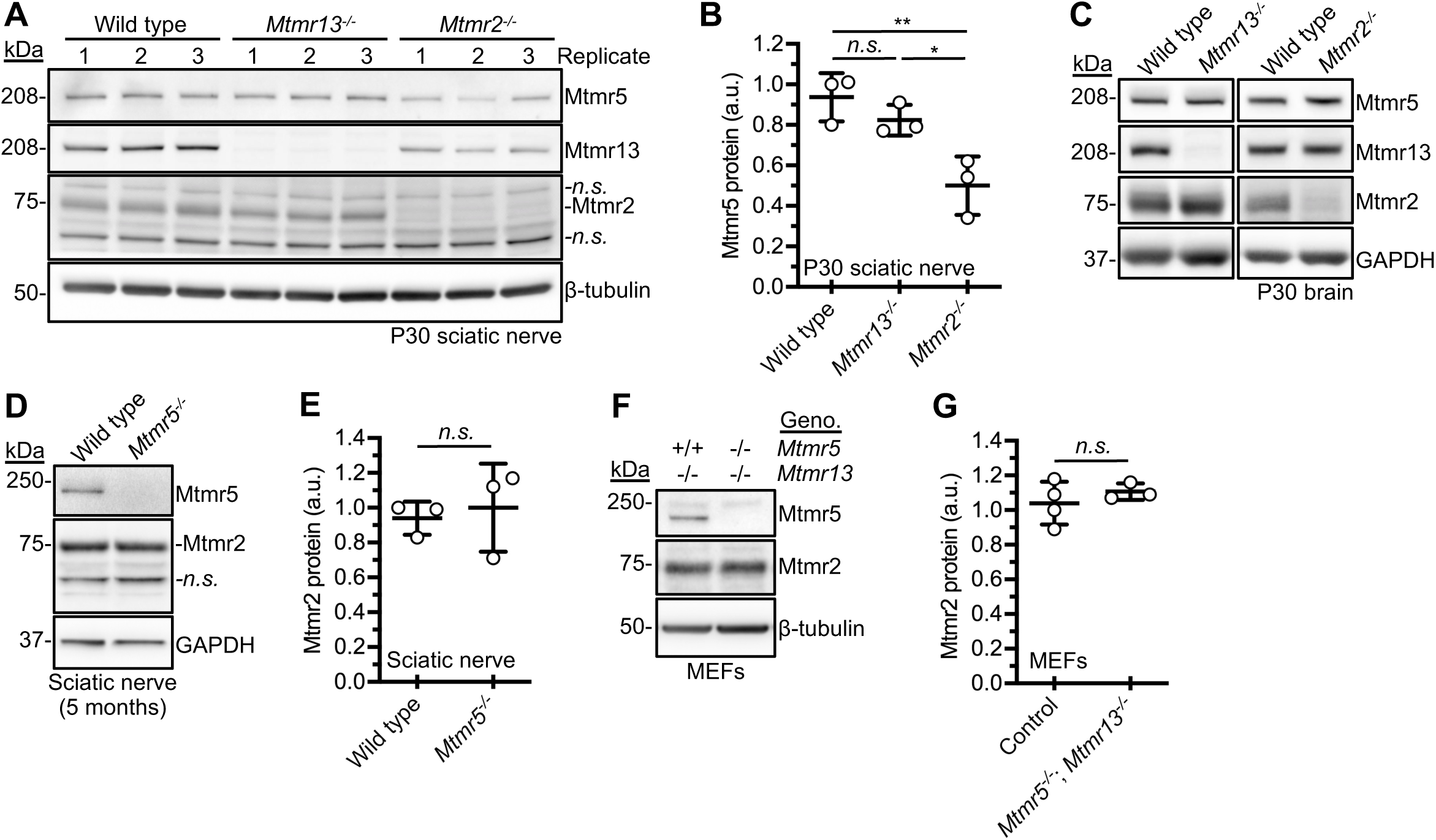
Relationships amongst CMT-linked myotubularin proteins assessed *in vivo.* (**A**) Sciatic nerve lysates were prepared from wild type, *Mtmr13^-/-^*, and *Mtmr2^-/-^* P30 mice. Pools 1, 2, and 3 denote independent (replicate) pools of nerves of the same genotype (4 nerves per lysate). Non-specific (*n.s.*) bands are indicated in Mtmr2 blots. (**B**) Relative quantification of Mtmr5 protein levels: wild type 0.937 ± 0.115, *Mtmr13^-/-^* 0.825 ± 0.072, *Mtmr2^-/-^* 0.499 ± 0.147 (arbitrary units, a.u.; mean ± SD; one-way ANOVA with post-hoc Tukey test; \**p* < 0.05; \*\**p* < 0.01; *n.s.,* not significant; n = 3 for each genotype). (**C**) Protein extracts from wild type, *Mtmr13 ^-/-^*, and *Mtmr2^-/-^* P30 brain lysates showed no significant differences in Mtmr5 protein levels. Representative data from one of three independent experiments are shown. (**D**) Sciatic nerve lysates prepared from wild type and *Mtmr5^-/-^* animals at age 5 months (2 nerves per lysate) were analyzed by immunoblotting. (**E**) Quantitation of (D), Mtmr2 protein: 0.937 ± 0.096 for wild type *vs.* 1.003 ± 0.253 for *Mtmr5^-/-^* nerves (*p* = 0.697; n = 3 for each genotype; two-sample Student’s t-tests; *n.s.,* not significant). (**F**) Protein extracts from control (*Mtmr5^+/+^*; *Mtmr13^-/-^*) and *Mtmr5^-/-^*; *Mtmr13^-/-^* MEF cultures were immunoblotted for Mtmr2. (**G**) Quantitation of F. Significant differences in Mtmr2 protein levels were not observed. Mtmr2 protein: 1.040 ± 0.122 for controls (*Mtmr5^+/+^*; *Mtmr13^-/-^* or *Mtmr5^+/-^*; *Mtmr13^-/-^*) *vs.* 1.104 ± 0.048 for *Mtmr5^-/-^*; *Mtmr13^-/-^*, respectively (mean ± SD*; p* = 0.258; n = 3-4 per genotype; two-sample Student’s t-tests; *n.s.,* not significant).

The stabilization of Mtmr5/13 by Mtmr2 was unique to the PNS; loss of Mtmr2 had no impact on the levels of Mtmr5 or Mtmr13 in brain extracts (Fig. 4C). In addition, loss of both Mtmr5 and Mtmr13 in *Mtmr5^-/-^*; *Mtmr13^-/-^* mouse embryonic fibroblasts (MEFs) did not impact Mtmr2 abundance (Fig. 4F, G). In summary, these data demonstrate that Mtmr2 is needed for wild type levels of both Mtmr5 and Mtmr13 protein abundance, specifically in the PNS, where CMT disease manifests.

### Reduced sciatic nerve axons and normal myelination in the absence of Mtmr5

In mice, the nerve pathology caused by the loss of Mtmr13 is well-characterized (33, 34). In contrast, the mouse nerve pathology associated with the loss of Mtmr5 is undefined. Loss-of-function mutations in *MTMR13/Mtmr13* cause distinct Schwann cell myelin abnormalities known as myelin outfoldings in both CMT patients and mouse models (33–35). The role of MTMR5 in the PNS remains unclear; significantly different pathology has been described in patient nerve biopsies (18, 27, 28). Through examination of *Mtmr5^-/-^* mouse nerves, we sought to determine whether Mtmr5 has a distinct or analogous role to that of Mtmr13 in the PNS. At 3 months, the morphology of *Mtmr5^-/-^* sciatic nerves appeared grossly normal (Fig. 5A-D). Loss of Mtmr5 did not cause CMT4B-like myelin outfoldings, indicating a pathology distinct from that caused by the loss of Mtmr13 (Fig. 5C-E). The absence of Mtmr5 did not alter myelin thickness; g-ratios were not significantly different between *Mtmr5^-/-^* and wild type fibers (Fig. 5F). However, *Mtmr5^-/-^* mice had significantly fewer total myelinated axons in sciatic nerves than wild type controls (Fig. 5G). The absence of Mtmr5 did not alter the diameter of myelinated axons, a feature of several mouse models of axonal CMT (Fig. 5H) (36, 37). These data demonstrate that Mtmr5 deficiency causes nerve pathology distinct from that caused by the absence of Mtmr13. Thus, despite sharing a common binding partner (Mtmr2), and possessing similar protein structures, the two pseudophosphatases play distinct roles in the murine PNS.

**Figure 5:**
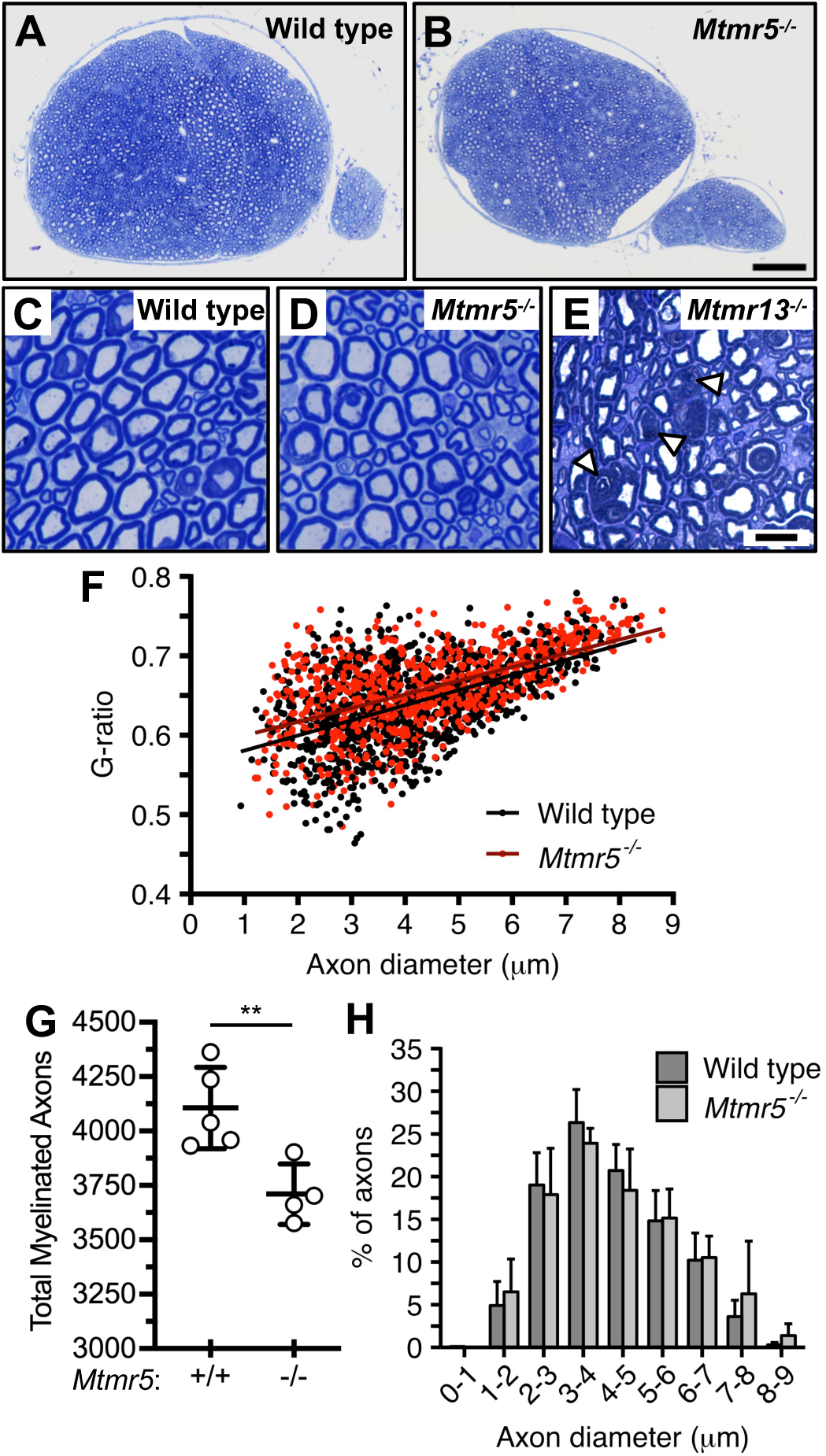
Sciatic nerve morphology in the absence of Mtmr5. (**A, B**) Comparable sciatic nerve cross-sections from wild type and *Mtmr5^-/-^* mice are shown (3 months of age). Scale bar: 100 μm. (**C, D**) Toluidine blue-stained mid-sciatic nerve myelin appeared normal in *Mtmr5^-/-^* mice; CMT4B-like myelin outfoldings were not observed. Scale bar: 10 μm. (**E**) A comparable image of an *Mtmr13^-/-^* mid-sciatic nerve stained with toluidine blue. Arrowheads indicate myelin outfoldings. (**F**) G-ratio analysis of myelin thickness in wild type and *Mtmr5^-/-^* nerves (best-fit line; N = 4-5 mice; 200 myelinated axons per mouse). (**G**) The total number of myelinated axons in sciatic nerves of *Mtmr5^-/-^* mice was significantly reduced at 3 months of age (mean ± SD; \*\**p* < 0.01; two-sample Student’s t-test; n = 4-5 mice per genotype). (**H**) Similar axon size distribution between wild type and *Mtmr5^-/-^* sciatic nerves (Two-sample Kolmogorov-Smirnov test; *p* = 0.16; wild type, N = 5 mice, n = 999 axons; *Mtmr5^-/-^*, N = 4 mice, n = 799 axons).

### Axonal sorting defects in the absence of Mtmr5

Incomplete radial sorting by immature Schwann cells can lead to the abnormal retention of large caliber axons in bundles, thereby reducing the total number of myelinated axon fibers in nerves (38, 39). To determine whether the absence of Mtmr5 caused abnormal Remak bundle development, we examined axon bundles of the sciatic nerve area using EM. Wild type and *Mtmr5^-/-^* sciatic nerves were characterized by determining the total number of axons per bundle, the proportion of bundled axons whose diameter exceeded 1 μm, and whether bundled axons were incompletely enclosed by Schwann cell processes.

In *Mtmr5^-/-^* nerves examined at 3 months, we observed a significant increase in the proportion of bundled axons that had a diameter of >1 μm (Fig. 6A-C). We also observed a significant increase in single unmyelinated axons, and a decrease in the percentage of bundles containing between 11-20 axons (Fig. 6D), suggesting delayed sorting of large axons from bundles, and abnormal retention of such axons in bundles. In addition, bundled axons of *Mtmr5^-/-^* nerves were significantly more likely to be incompletely enwrapped by Schwann cell processes, as evidenced by direct contact of axons with basal lamina (Fig. 6E-H).

**Figure 6:**
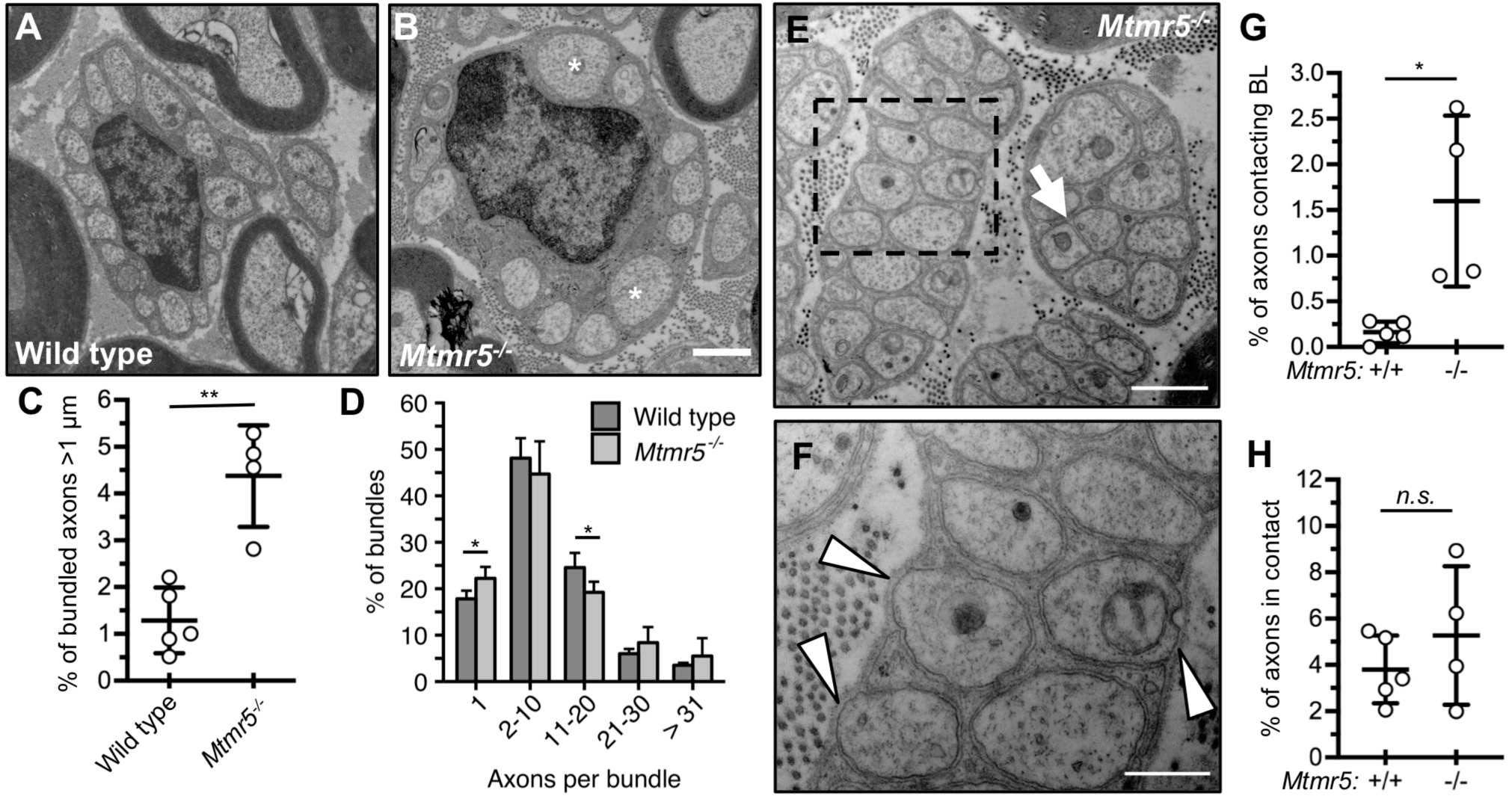
The absence of Mtmr5 alters axon radial sorting and ensheathment. (**A, B**) EM images of representative axon bundles in sciatic nerves of wild type and *Mtmr5^-/-^* mice at 3 months. Asterisks indicate axons >1 μm in diameter. Scale bar: 1 μm. (**C**) The percentage of bundled axons greater than 1 μm in diameter was significantly increased in *Mtmr5^-/-^* sciatic nerves at 3 months (1.29 ± 0.70% for wild type *vs.* 4.37 ± 1.09% for *Mtmr5^-/-^*, respectively; n = 4-5 mice per genotype). (**D**) Number of axons per bundle expressed as a percentage. Categories are distinguished by the number of ensheathed axons. (**E**) EM of two bundles from a transverse *Mtmr5^-/-^* sciatic nerve section. The arrow indicates an area without Schwann cell cytoplasm between axons. Scale bar: 1 μm. (**F**) Higher magnification image of the section boxed in image (E). Arrowheads identify points of direct contact between the basal lamina (BL) and axons. Scale bar: 500 nm. (**G, H**) Axons lacking full Schwann cell ensheathment were counted and expressed as a percentage of the total bundled axons per mouse. (**G**) Quantification of the percentage of axons abnormally contacting the Schwann cell BL. (**H**) Quantification of the percentage of axons in direct contact with another axon (mean ± SD; Student’s t-test; \**p* ≤ 0.05; **p ≤ 0.01; n = 5 wild type and n = 4 *Mtmr5^-/-^* mice).

In aggregate, our findings indicate that Mtmr5 regulates late-stage radial sorting of large caliber axons. We propose that, after the completion of radial sorting, Mtmr5 is dispensable for subsequent myelination.

### Loss of both Mtmr5 and Mtmr13 is lethal in mice

Mtmr5 and Mtmr13 share 59% protein identity and are the only two proteins in the genome that share their unique set of protein domains (Fig. 1) (40). To assess whether these proteins might have partially redundant functions, we generated *Mtmr5^-/-^*; *Mtmr13^-/-^* double knockout (dKO) mice. dKO animals were non-viable, dying during late gestation or within a few hours of birth (Fig. 7). *Mtmr5^-/-^; Mtmr13^-/-^* embryos were observed at the expected Mendelian frequencies at embryonic (E) days 13-15, but were smaller than littermate controls; overt morphological defects were not apparent (Fig. 7, Supp. Fig. S4, and *data not shown*). In summary, the absence of both Mtmr5 and Mtmr13 leads to physiological deficits that preclude post-natal survival.

**Figure 7:**
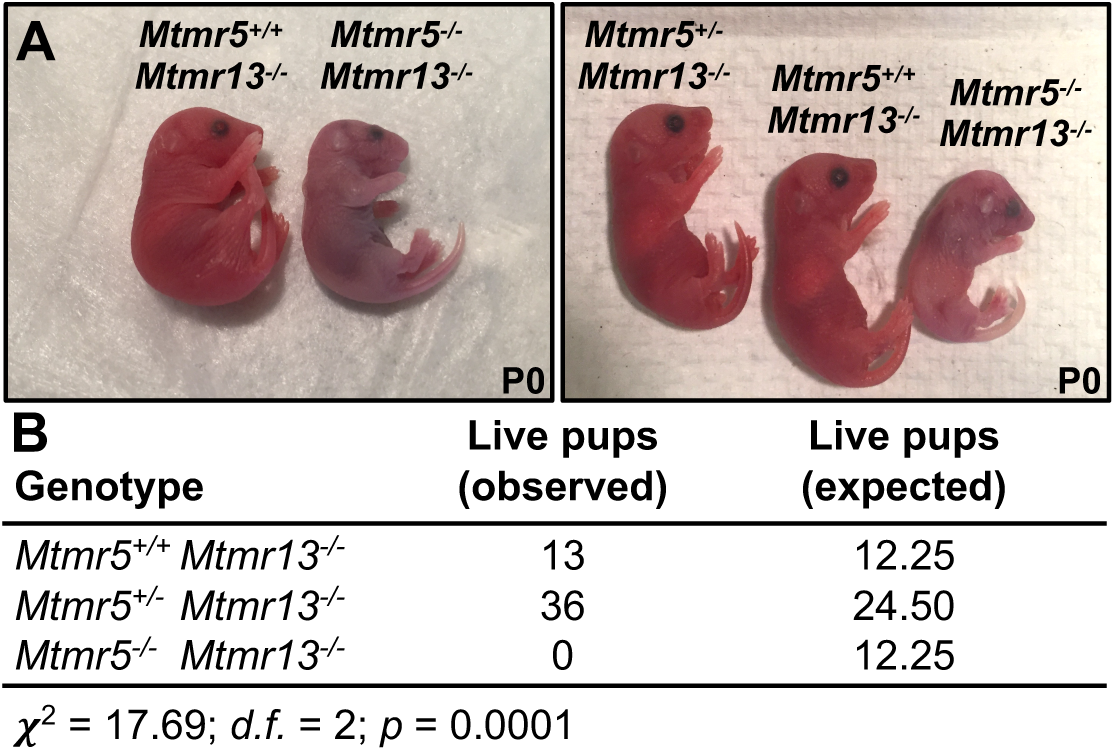
*Mtmr5-Mtmr13* double-knockout mice are not viable beyond birth. (**A**) Images of *Mtmr5^-/-^*; *Mtmr13^-/-^* mice found dead at P0, alongside littermate controls. (**B**) *Mtmr5^-/-^*; *Mtmr13^-/-^* double knockout animals were not represented at the expected ratios (Chi-squared (χ^2^) 17.69; *p* = 0.0001; n = 49 total live pups).

Mtmr5 and Mtmr13 have critical roles in the PNS, and both are proposed to activate the early endosomal GTPase Rab21, a regulator of neurite growth and axon guidance (41, 42). Accordingly, we assessed whether the combined loss of both proteins might alter axon guidance and/or peripheral nerve development during embryogenesis. We found that nerve branches in dKO mouse pinna, a region where axon guidance is well defined, appeared morphologically similar to those of controls (Supp. Fig. S4). Thus, a strict requirement for Mtmr5 or Mtmr13 for axon guidance appears unlikely.

### Myotubularin expression in peripheral nerves and Schwann cells

We considered whether the phenotypic differences between *Mtmr5^-/-^* and *Mtmr13^-/-^* mice might correlate with the expression levels of the two proteins during PNS development. Mtmr2, Mtmr5, and Mtmr13 protein levels in sciatic nerves were therefore evaluated at distinct time points during mouse development (Fig. 8A, B). Mtmr5 levels were highest between P0 and P7, but decreased sharply by P21 (Fig. 8A, B). In contrast, Mtmr13 levels increased between P0 and P7, peaked at P21, and decreased moderately thereafter (Fig. 8A, B). Mtmr2 protein levels remained relatively constant throughout sciatic nerve development (Fig. 8A, B). Thus, Mtmr5 levels were highest during late stage radial sorting (E17-P10), consistent with our morphological findings indicating a role for the protein in this process. In contrast, Mtmr13 protein expression correlated with initial myelination (P0-P14), and, to a lesser degree, with myelin maintenance in adulthood.

**Figure 8:**
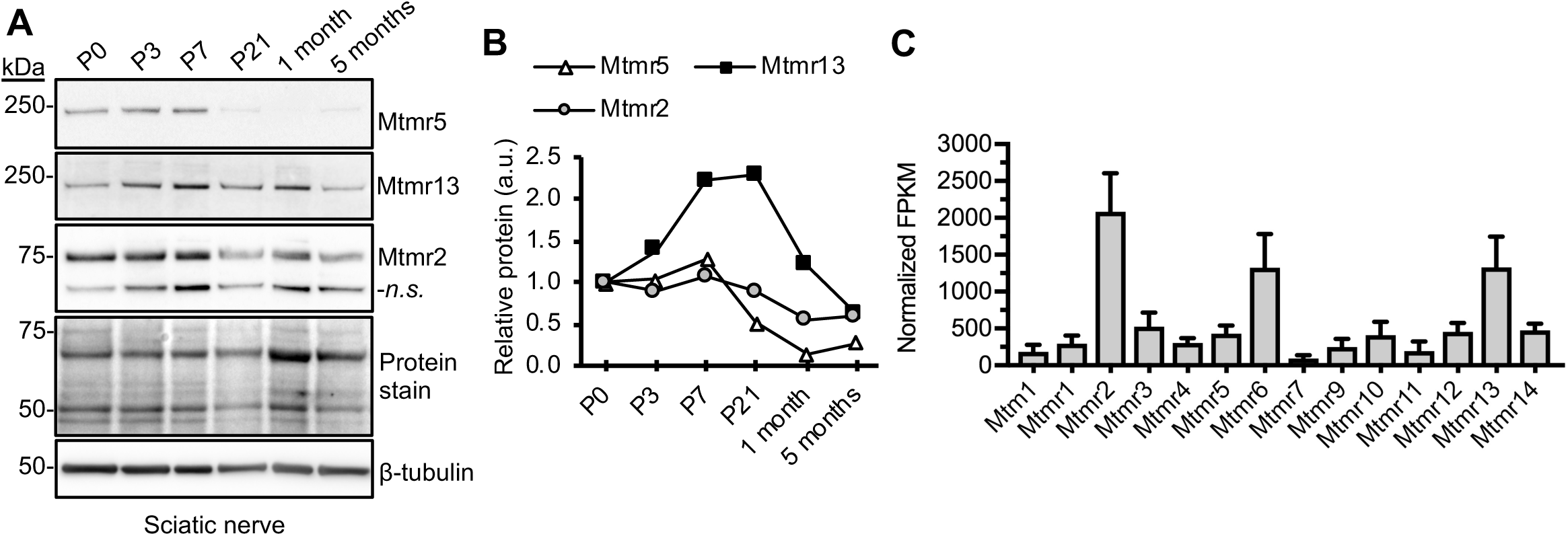
Myotubularin protein expression in sciatic nerve and Schwann cells. (**A**) Immunoblot of Mtmr5, Mtmr13, and Mtmr2 proteins in wild type mouse sciatic nerve extracts at P0, P3, P7, P21, 1 month, and 5 months of age. A β-tubulin immunoblot and a total protein stain are shown as loading controls. (**B**) Relative protein abundance of Mtmr5, Mtmr13, and Mtmr2 at each age, normalized to the total protein stain. For (A) and (B), the data from a single experiment are shown. (**C**) Relative mRNA expression of all 14 myotubularins in Schwann cells isolated from wild type adult mouse sciatic nerves, presented as RNAseq-derived FPKM. The graph was generated using the publicly available *Clements, M.P. et al.* dataset (GSE103039) (43). Abbreviation: FPKM, Fragments Per Kilobase of transcript per Million mapped reads.

**Figure 9:**
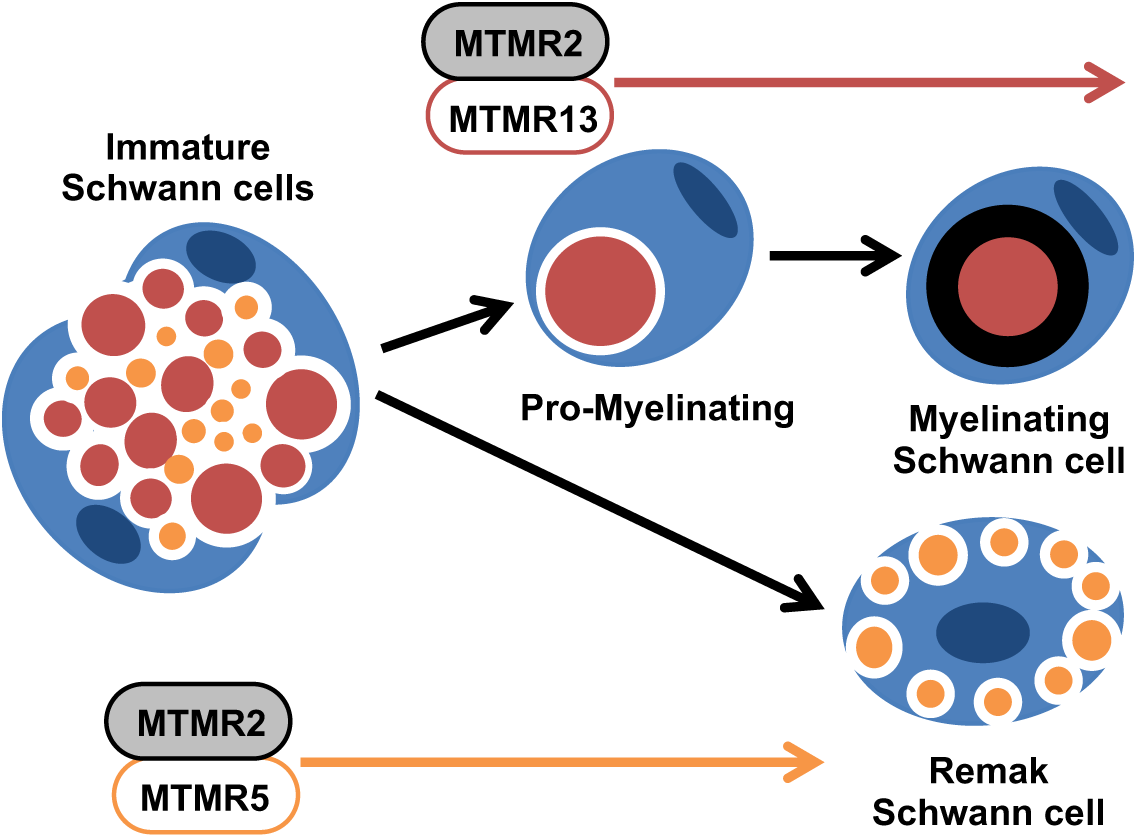
Proposed model for the respective roles of Mtmr5 and Mtmr13 in the PNS. We propose that two distinct myotubularin protein complexes form in the PNS: an Mtmr2-Mtmr5 complex and an Mtmr2-Mtmr13 complex. An Mtmr2-Mtmr5 protein complex may positively regulate late stage axon radial sorting (E17 to P10). Subsequently, an Mtmr2-Mtmr13 protein complex may function to prevent abnormal myelin formations, once large axons have formed 1:1 relationships with promyelinating Schwann cells (P0 to adulthood). Differences in the timing of Mtmr5 and Mtmr13 expression may partially explain the impact of the loss of the two proteins on axon sorting and myelination, respectively.

We also evaluated the mRNA levels of all 14 myotubularin family members in adult mouse Schwann cells by analyzing a publicly available RNA-Seq dataset (43). This analysis indicated that *Mtmr2* and *Mtmr13* are expressed at significantly higher levels than nearly all other *Mtmr* mRNAs in the Schwann cells of adult nerves (Fig. 8C). *Mtmr5* mRNA was present at low levels, consistent with our protein expression findings (Fig. 8A-C). The observed temporal differences in the expression of Mtmr5 and Mtmr13 correlate with the distinct PNS phenotypes of *Mtmr5^-/-^* and *Mtmr13^-/-^* mice.

## DISCUSSION

### Mtmr5-deficient mice as a model CMT disease

Loss of Mtmr5 in mice caused a ∼10% reduction in the number of myelinated axons in the sciatic nerve due to radial sorting defects. Several patient families with loss-of-function mutations in *MTMR5* show moderate depletion of myelinated axons without defects in myelin structure, suggesting primary axonal neuropathy (28, 29). Given the findings presented here, we propose that abnormal axon radial sorting be considered as a potential primary defect in CMT associated with *MTMR5* loss-of-function. The effects of *MTMR5* loss-of-function may be exacerbated in humans relative to mice, potentially explaining the greater reductions in myelinated axon numbers observed in patients (28, 29). Although the precise cause of this feature is unclear, several mouse models of axonal CMT have shown phenotypes milder than their corresponding human conditions (36, 44–48). Incomplete radial sorting can be caused by defects in both axons and Schwann cells (1, 49). Accordingly, future studies employing conditional *Mtmr5* knockout alleles will be needed to determine if Mtmr5’s role is restricted to axons or Schwann cells.

In the *Mtmr5^-/-^* PNS, myelin was of normal thickness and structure, and notably lacked the distinctive myelin outfoldings caused by the absence of Mtmr2 or Mtmr13. Nerve biopsies from a family of CMT patients with compound heterozygous mutations in *MTMR5* (M417V and T1590A) showed CMT4B-like myelin outfoldings (18) (Fig. 1). Our study demonstrates that a *null*-type mutation in *Mtmr5* does not provoke CMT4B-like myelin outfoldings in mice. Therefore, we speculate that the M417V and T1590A mutations in MTMR5 may trigger CMT in a manner distinct from that caused by MTMR5 elimination (18). One possible mechanism for such an affect might be that mutations M417V and/or T1590A alter MTMR5 in a manner that changes the pseudophosphatase’s interaction with its binding partner MTMR2. If a mutated MTMR5 protein were to impair MTMR2 function, myelin outfoldings might be predicted to result.

### Mtmr5 and Mtmr13 form distinct protein complexes with Mtmr2 in the PNS

Both Mtmr5 and Mtmr13 levels were enhanced when the proteins were expressed with their binding partner Mtmr2; this effect was observed both *in vitro* and in the PNS. We showed that Mtmr5 and Mtmr13 form a very weak interaction that was independent of Mtmr2. If extant, an Mtmr5-Mtmr13 complex would be incapable of dephosphorylating PI3P or PI(3, 5)P_2_, key regulators of trafficking through early endosomes and lysosomes. Therefore, we hypothesize that two distinct myotubularin protein complexes form in the PNS: an Mtmr2-Mtmr5 complex and an Mtmr2-Mtmr13 complex. These two complexes would have similar capacity to regulate endosomal trafficking by activating Rab GTPases and dephosphorylating PIs. In tissues outside the PNS, the Mtmr2-Mtmr5 and Mtmr2-Mtmr13 complexes likely have overlapping or redundant functions, as loss of both Mtmr5 and Mtmr13 caused perinatal lethality.

We show here that, despite their redundant role(s) during embryonic development, Mtmr5 and Mtmr13 play distinct biological roles in peripheral nerves. The differing expression patterns of these two homologous proteins during nerve development may at least partially explain the stark difference between the *Mtmr5^-/-^* and *Mtmr13^-/-^* nerve phenotype. Mtmr5 levels were high during axon radial sorting, and decreased sharply thereafter. In contrast, Mtmr13 levels increased notably between birth and P21, a period coinciding with initial myelination, and were only moderately decreased in adult nerves. Mtmr2 protein levels in sciatic nerves changed little from birth to adulthood. Accordingly, we propose that Mtmr2 and Mtmr5 function together in immature Schwann cells to control which axons are sorted for myelination. Once large axons form 1:1 relationships with Schwann cells, an Mtmr2-Mtmr13 complex may function to ensure orderly myelin wrapping and maintain myelin homeostasis in adulthood. The lack of Mtmr5 protein expression after P7 may explain why the loss of Mtmr5 did not lead to reduced Mtmr2 protein in adult nerves; reduced Mtmr2 levels were observed in *Mtmr13^-/-^* mice (22). Likewise, the uniform expression of Mtmr2 during all stages of nerve development may explain why the absence of this protein leads to decreases in both Mtmr5 and Mtmr13 levels in sciatic nerves. In summary, we demonstrate here that, in most tissues, either Mtmr5 or Mtmr13 protein is sufficient for normal cell function. In contrast, the PNS may require both proteins because of temporal differences in *Mtmr5* and *Mtmr13* expression.

### Control of endosome trafficking by Mtmr5 and Mtmr13 during axon sorting and myelination

A majority of the molecularly-defined forms of demyelinating CMT are caused by mutations in endosomal regulatory proteins (12). Schwann cells select large axons for myelination by integrating signals from ErbB2/3 tyrosine kinase receptors on their cell surfaces (50–52). Abnormal trafficking of the ErbB2/3 receptor complex in Schwann cells has been proposed as a common mechanism linking multiple demyelinating subtypes of CMT (16). Our previous work, and that of others has suggested that appropriate regulation of endosomal PI3P levels in Schwann cells is critical to controlling ErbB2/3 trafficking, and the associated downstream signaling (9, 24). Specifically, the loss of Mtmr2 has been shown to increase ErbB2/3 activation during initial myelination (24). As both Mtmr5 and Mtmr13 physically associate with Mtmr2, we speculate that loss of either pseudophosphatase may also lead to increased ErbB2/3 activation, likely by altering receptor trafficking through endosomes. Further investigations focused on Mtmr5 and Mtmr13 may serve to clarify the dual role of ErbB2/3 in controlling axon sorting and myelination (53, 54).

We speculate that Mtmr5 and Mtmr13 might also influence the trafficking of β1-integrin, which binds to laminin 211 in the Schwann cell basal lamina. The trafficking of β1-integrin through early endosomes is controlled by Rab21, which the DENN domain of Mtmr5/13 has been shown to activate (42). β1-integrin is required for the extension and maintenance of Schwann cell processes around axons; loss of this surface receptor leads to radial sorting defects and delayed myelination in mice (55). In *Mtmr5^-/-^* nerves, we observed an increase in the proportion of unmyelinated axons that were incompletely ensheathed by Schwann cells. We speculate that this defect might be caused by a reduction in β1-integrin on the surface of *Mtmr5^-/-^* Schwann cells, a result of reduced recycling of the receptor from early endosomes. An analogous defect in β1-integrin trafficking has been demonstrated in Mtmr5/13-deficient *Drosophila* cells (42). While ErbB2/3 and β1-integrin may presently be the best candidates for the signaling receptors influenced by Mtmr5/13-mediated endosomal trafficking, other yet to be identified surface proteins may also be regulated by these pseudophosphatases. Understanding how Mtmr5 and Mtmr13 regulate the trafficking of receptors involved in mediating axo-glial interactions and promyelinating signaling will likely be critical to discerning how the loss of these proteins leads to CMT.

In conclusion, we have shown, for the first time, that Mtmr5 positively regulates axon radial sorting in mouse peripheral nerves. This work demonstrates that Mtmr5 plays a biological role district from that of the homologous Mtmr13 protein, the loss of which causes demyelinating CMT4B2 in humans. Our study may inform investigations of the forms of CMT that result from loss-of-function mutations in *MTMR5,* and may aid in the definition of the cellular mechanisms that underpin this condition.

## MATERIALS AND METHODS

### Mice

All animal work was approved by and conformed to the standards of the Oregon Health & Science University Animal Care and Use Committee. *Mtmr2^-/-^* and *Mtmr13^-/-^* mice have been previously described (22, 33). An *Mtmr5* deletion allele was generated via CRISPR/Cas9 mutagenesis. To do so, two guide RNAs were designed against exon 1 and exon 25 of the mouse *Mtmr5* gene using the online tool CRISPR Design. The following guide RNAs (gRNAs) were selected for their minimal off-target affects and potential to generate a large deletion beginning after the start-site (exon 1: GCTCGCGGACTACTTCGTGC) and ending after a putative internal initiation site (exon 25: CGTAAGAAGTATAACCCCCC). These *Mtmr5-*specific gRNA sequences were ligated into the *BbsI* restriction endonuclease site of px330 (addgene plasmid #42230) (56). Plasmids containing gRNAs plus Cas9 endonuclease were injected into the pronuclei of C57BL/6NJ mouse embryos. The embryo donors were oviduct transferred into CB6F1/J pseudopregnent recipient dams. *Mtmr5* founder mice were screen by PCR and Sanger sequencing to identify putative mutations in exon 1, exon 25, and large deletions between exon 1 and exon 25 using the following primers: forward exon 1 primer AMO-49 5’ CATGCGGAGTGGCCCAAT 3’, reverse exon 1 primer AMO-50 5’ GGATGTTTCTTACACAGGCCATGT 3’, forward exon 25 primer AMO-51 5’ CACGGGTTACCAAGGACAAGG 3’, reverse exon 25 primer AMO-56 5’ GTCAACTCTGATAGCGAGCACAG 3’. Mutations were confirmed by TOPO TA (ThermoFisher) cloning and Sanger sequencing. Mice were breed to homozygosity and western blots were run to confirm Mtmr5 protein loss in brain and sciatic nerve tissue. To test male sterility, six *Mtmr5^-/-^* male mice were allowed to breed with C57BL/6 females for at least 21 days. Bred females were monitored for pregnancy and births for an additional three to four weeks.

*Mtmr5^-/-^*; *Mtmr13^-/-^* (double knockout) mice were generated by crossing the *Mtmr5* CRISPR founder male with a *Mtmr13^-/-^* female. *Mtmr5^+/-^*; *Mtmr13^+/-^* progeny were screened by PCR and Sanger sequencing for the *Mtmr5* large deletion using the following primers: AMO-49 5’ CATGCGGAGTGGCCCAAT 3’ and AMO-56 5’ GTCAACTCTGATAGCGAGCACAG 3’. The *Mtmr5^+/-^*; *Mtmr13^+/-^* mice were crossed with *Mtmr13^-/-^* mice to generate *Mtmr5^+/-^*; *Mtmr13^-/-^* progeny. The *Mtmr5^+/-^*; *Mtmr13^-/-^* mice were interbred and their pups were genotyped at P0 to screen for dKO animals. 49 pups from 13 litters were genotyped to assess the viability of Mtmr5 and Mtmr13 loss.

### Plasmid Constructs

Mammalian expression vectors for human FLAG-MTMR2 (26), human EGFP-MTMR2 (57), human EGFP-MTMR13, and myc-MTMR13 mutant constructs (32) have been described. A human FLAG-MTMR5 expression vector (26) was modified to eliminate a single glutamine insertion between amino acid 93 and 94 and a K592M mutations. To generate the wild type construct human FLAG-MTMR5-AM1, transcript variant 1 from NM-002972.3 was amplified using oligonucleotides: 5’ GTTTAAACTTAAGCTTATGGCGCGGCTCGCGGAC 3’ and 5’ CCTGATGTCCGGCGGGTACCG 3’. The resulting PCR product was cloned into the previously described FLAG-MTMR5 construct, which had been digested with *KpnI* and *HindIII*, using In-Fusion (Takara Bio).

### Cell Culture and Immunoprecipitation

HEK293 cells were cultured in Dulbecco’s Modified Eagle’s Medium (DMEM; Gibco 11995-065), that was supplemented with 10% fetal bovine serum (FBS) and 1% penicillin/streptomycin. Cells were between 70-80% confluent at the time of transfection. A 60 mm dish was transfected with 2.5 μg of plasmid DNA, 7.5 μL X-tremeGENE9 (Roche), and 250 μL of Opti-MEM (Gibco) according to manufacturer’s instructions. Forty-eight hours after transfection, cells were washed with 1x phosphate-buffered saline (PBS) and lysed in 430 μL of ice-cold lysis buffer (120 mM NaCl, 50 mM Tris [pH 8.0], 0.5% Triton X-100, 100 mM NaF, 1 mM ortho-vanadate, 2 mM EDTA, and a protease inhibitor cocktail [Roche 11836153001]). Lysates were vortexed and cleared by centrifugation (15,000 x g for 15 min at 4°C). Myc-tagged proteins were immunoprecipitated using anti-c-Myc monoclonal antibody supernatant (9E10, DSHB) and protein A-agarose (Invitrogen). Flag-tagged proteins were immunoprecipitated using anti-Flag-M2 affinity gel (Sigma). Immunoprecipitates were incubated for at least 2 h at 4°C, washed twice in 1 ml of lysis buffer, three times in 1 ml of lysis buffer containing 0.5 M NaCl, and once again in 1 ml of lysis buffer. Immunoprecipitates and lysates were suspended in 1x NuPAGE LDS sample buffer containing 10 mM DTT. Immunoprecipitates and lysates were resolved in 4-12% NuPAGE BisTris gels (Life Technologies) and transferred to polyvinylidene difluoride membranes for immunoblotting.

### Immunoblotting

Immunoblotting of sciatic nerve, brain, HEK293 cell, and mouse embryonic fibroblast (MEF) extracts was performed as previously described (22). Rabbit antibodies anti-MTMR13 (116-AN) and anti-MTMR2 (119-AN) have been described (22). Mouse anti-FLAG (M2) and anti-c-myc (9E10) were from Sigma-Aldrich and Roche Applied Science, respectively. Mouse anti-Mtmr5 (B-9) was from Santa Cruz Biotechnology. Mouse anti-GFP antibody (N86/8) was from NeuroMab. Mouse anti-β tubulin (E7) and anti-GAPDH (MAB374) were from DSHB and Millipore, respectively. For each P0, P3, and P7 lysate, nerves from three mice were pooled (6 nerves per lysate). For P21 and one-month mouse extracts, nerves from two mice (4 nerves total) were pooled to generate each independent extract. Extracts from mice 3 months and older were generated from a single animal (2 nerves per lysate). For each immunoblot, 8-25 μm of protein per lane was resolved in 4-12% NuPAGE Bis-Tris gels (Invitrogen) and chemiluminescent quantitation was performed as previously described (23).

### Sciatic Nerve Morphology

Mice were perfused with Karnovsky’s EM fixative (4% paraformaldehyde, 2% glutaraldehyde; 0.1 M sodium cacodylate, pH 7.4). Sciatic nerves were dissected and further fixed for at least 24 hours at 4°C. Nerves were washed three times in 0.1 M sodium cacodylate (pH 7.4) and three times in 0.1 M sodium phosphate buffer (PB; pH 7.4) at room temperature (RT). Nerves were subsequently post-fixed and stained in 2% osmium tetraoxide in 0.1 M PB buffer (pH 7.4) for 1 hr. The nerves were then washed three times in 0.1 M PB at RT and underwent a series of ethanol dehydrations (25%, 50%, 70%, 80%, and 95%; 5 min each). A final dehydration was performed by incubating samples twice in 100% EtOH (10 min each), followed by two incubations in propylene oxide (10 min each). Nerves were infiltrated overnight in a 1:1 propylene oxide:Embed 812 resin mixture, and subsequently in freshly-prepared 100% Embed 812 resin. Nerves were embedded in a 60°C oven for 48 h. The distal end was positioned toward the beveled end of the mold to orient the nerves in the same direction.

Toluidine blue-stained semi-thin (200-500 μm) plastic cross-sections were prepared from the mid-sciatic nerve. Four non-overlapping images of nerve sections were acquired at 63x on a Zeiss Axio Imager M2 ApoTome microscope with an AxioCam 512 color camera. The tile feature in Zeiss Zen 2 (blue edition) software was used to capture the entire transverse fascicular area (TFA) of each nerve. To obtain total axon counts, tiled 63x Toluidine blue-stained images were used. All myelinated axons in the TFA were marked and counted using Cell Counter plugin for Fiji (58). G-ratio was determined using the G-ratio plugin for Fiji. 200 myelinated axons were randomly selected across 4 non-overlapping 63x Toluidine blue images of the TFA. To obtain an area-based G-ratio measurement, the area of the axon was divided by the area of the myelinated fiber. This measurement was then used to derive a diameter-based G-ratio measurement.

EM images were obtained using a FEI Tecnai G2 microscope operating at 80 kV and an Advanced Microscopy Techniques camera. To analyze Remak bundles, 2900x EM images were used. All intact Remak axons in the large fascicle of each nerve were analyzed. Each Remak axon was counted, and its area was measured. Any axon with an area greater than 0.79 μm^2^ (1 μm in diameter) was considered large. 2900x EM images were also used to count abnormally ensheathed axons. At least 42% of the nerve TFA was exhaustively analyzed for each mouse.

### Immunofluorescence

Whole-mount immunofluorescence of E13 mouse embryos with neurofilament medium chain (2H3) antibody was adapted from a published protocol (59). Individual embryos were stained in microcentrifuge tubes at RT, unless otherwise stated. Embryos were separated from placenta and amnion in 1x PBS, and 3 mm of the tail was removed for use in genotyping. Embryos were fixed overnight at 4°C in 4% paraformaldehyde in 1x PBS solution (pH 7.4). After fixation, the embryos were washed with 1x PBS twice for 10 minutes. Embryos then underwent a series of methanol (MeOH) dehydrations (1 hr in 50% MeOH in 1x PBS, 2 hr 80% MeOH in 1x PBS, overnight in 100% MeOH). Embryos were bleached in 3% H_2_O_2_, 70% MeOH, 20% DMSO overnight, and washed five times at 45 minute intervals in 1x TNT (10 mM Tris, 154 mM NaCl, 0.1% Triton X-100). Bleached embryos were next incubated in a primary antibody solution (1x TNT, 0.02% sodium azide, 4% nonfat milk, 5% DMSO, 2% normal goat serum [NGS], 0.37 μg/mL 2H3 antibody) for 2-3 days. Embryos were washed five times in 1x TNT (45 min each), and subjected to a 2-day incubation in secondary antibody solution (1x TNT, 4% bovine serum albumin, 0.02% sodium azide, 5% DMSO, 2% NGS, 6.4 μg/mL Cy3 goat-anti-mouse (Jackson ImmunoResearch, 115-175-205)). In all subsequent steps, the embryos were protected from light. Embryos are washed four times in 1x TBS (pH 7.4) for 45 minutes, then overnight in the same solution, and dehydrated using a MeOH series: 1 hour in 50% MeOH (1x PBS), 2 hours in 80% MeOH (1x PBS), and overnight in 100% MeOH. The samples were cleared in 1:2 BABB solution (Benzyl:Benzyl benzoate). Whole embryo images were taken on Zeiss AxioCam MR microscope at 1x magnification prior to the 1:2 BABB clearing step. Images of nerve branches were taken on a Zeiss ApoTome microscope at 5x magnification in 1:2 BABB solution.

### Statistics

Statistical analyses of immunoblots, weight, and morphology data were performed using a one-way ANOVA followed by a post-hoc Tukey’s test when more than two groups were compared. A Student’s t-test was used when two groups were compared. A chi-squared (χ^2^) test was performed to evaluate the significance of divergences from the expected frequencies of specific genotypes. The nonparametric Kolmogorov-Smirnov test (K-S test) was used for comparing the distribution of axon size. A probability value of ≤ 0.05 was used for assigning statistical significance. χ^2^ and K-S tests were performed using R version 3.5.1. The Prism software package (GraphPad) was used to perform ANOVA, Tukey multiple comparisons, and Student’s t-tests.

## Acknowledgements

The authors wish to thank the OHSU Transgenic Mouse Models Shared Resource Core. The authors also wish to thank Matthew Pomaville and Kevin Wright for expert advice on mouse whole embryo immunofluorescence. The EM microscope was purchased through a Murdock Charitable Trust grant to Sue Aicher. The authors would like to thank Ben Emery and Kelly Monk for constructive criticism and substantial feedback on the manuscript. The E7 (β-tubulin) monoclonal antibody was developed by Dr. Michael Klymkowsky and obtained from Developmental Studies Hybridoma Bank (DSHB). The monoclonal antibody 2H3 (NF-M), developed by Drs. T.M. Jessell and J. Dodd, was obtained from the DSHB developed under the National Institutes of Health-National Institute of Child Health & Human Development and maintained by The University of Iowa, Department of Biology, Iowa City, IA 52242.

