## Supplemental Figures for "Distinct roles for the Charcot-Marie-Tooth disease-causing endosomal regulators Mtmr5 and Mtmr13 in axon radial sorting and Schwann cell myelination"

### Supplemental Figure S1 – Mammel *et al.*

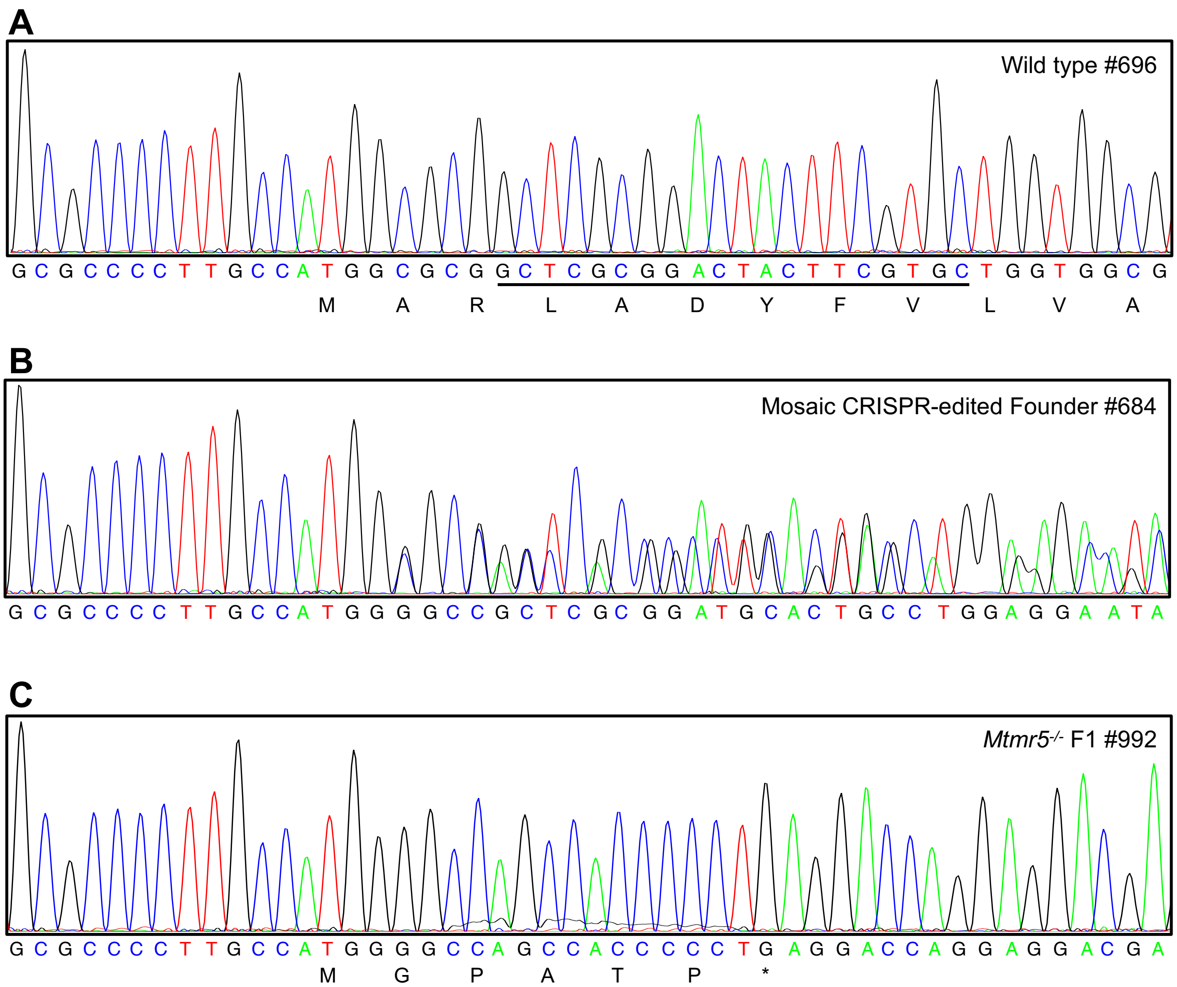

### Supplemental Figure S2 – Mammel *et al.*

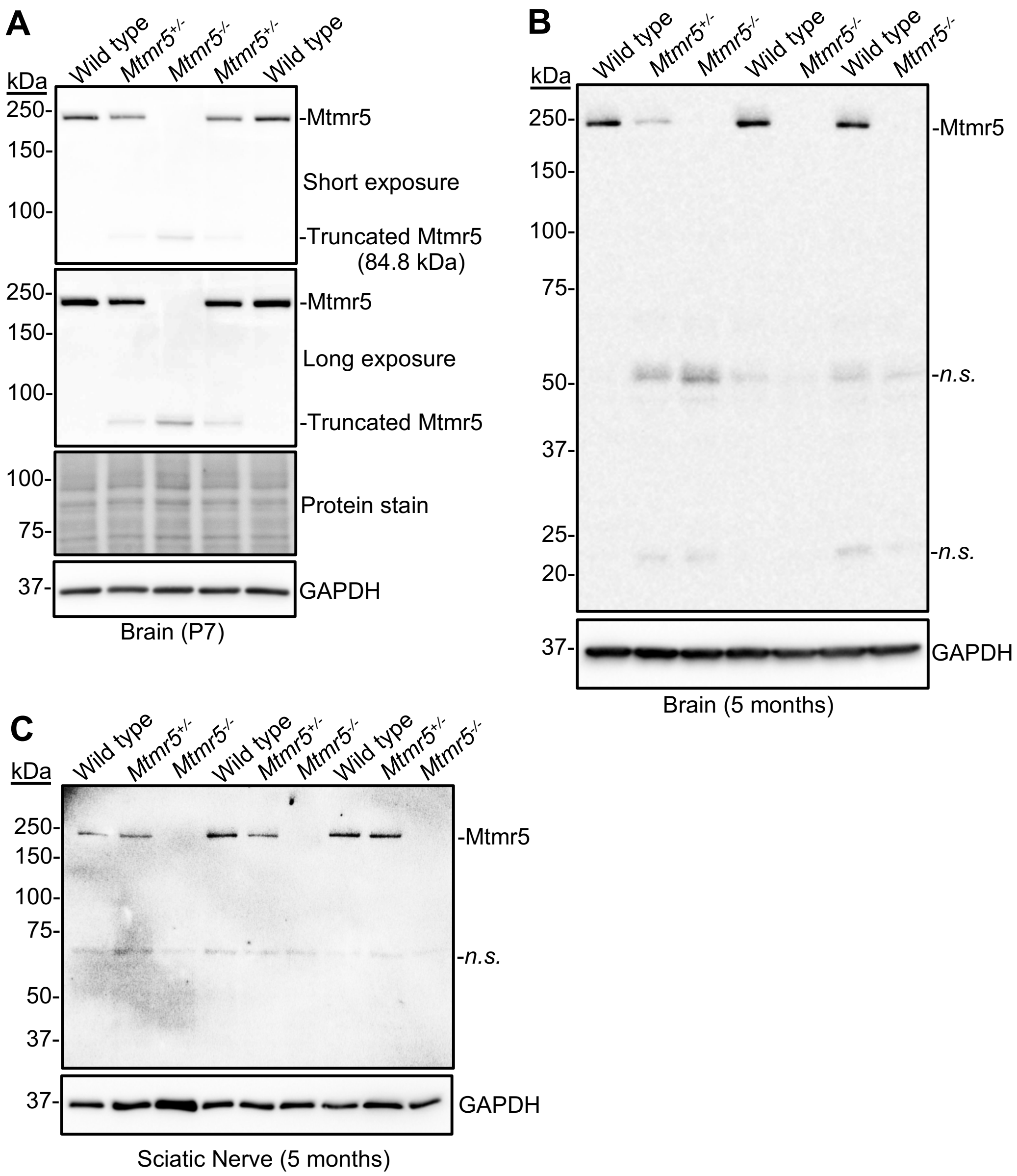

### Supplemental Figure S3 – Mammel *et al.*

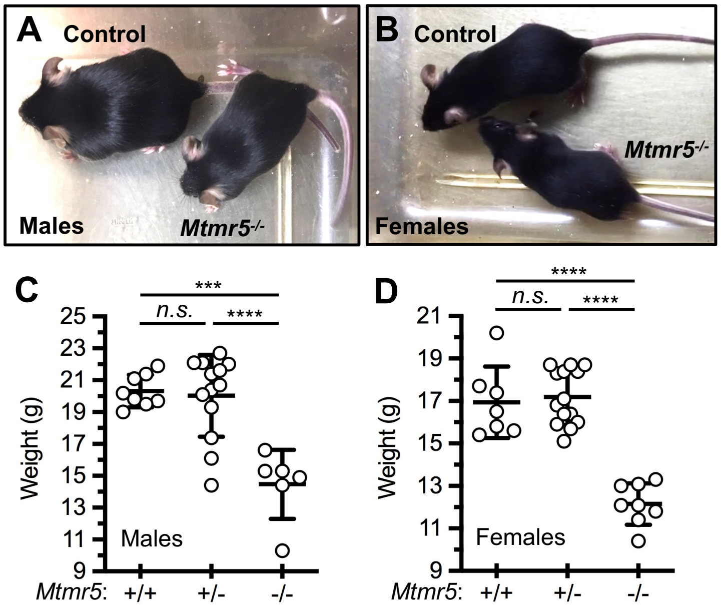

### Supplemental Figure S4 – Mammel *et al.*

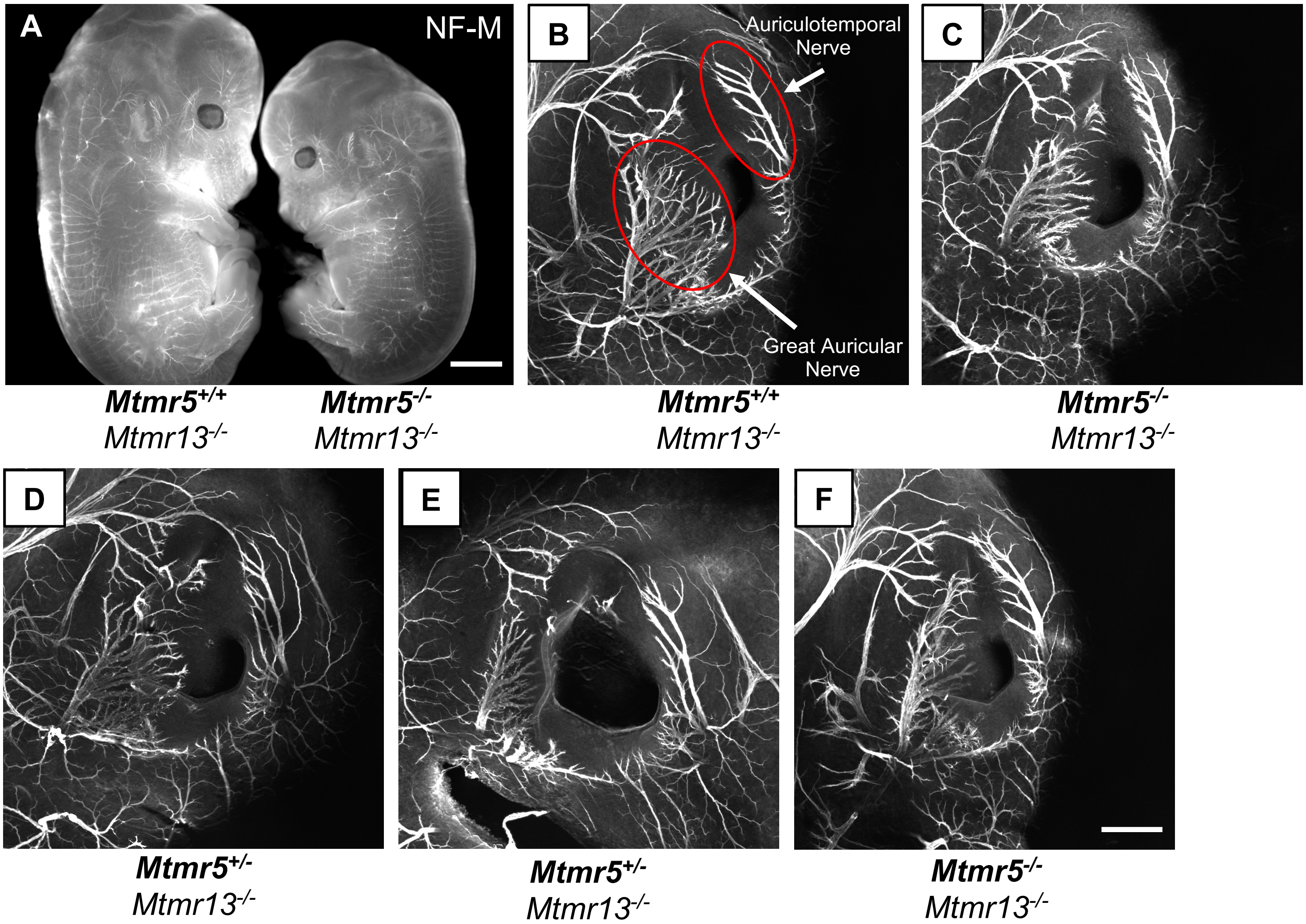

**SUPPLEMENTAL FIGURE LEGENDS – Mammel *et al.***

**Supplemental Figure S1:** DNA sequencing of *Mtmr5* deletion alleles generated via CRISPR/Cas9 gene editing. Representative chromatograms, obtained by Sanger sequencing of genomic PCR fragments containing the *Mtmr5* exon 1 gRNA target sequence, are shown. (**A**) Wild type mouse #696 chromatogram with gRNA sequence underlined and coding frame indicated. (**B**) CRISPR-Cas9 founder mouse #684 chromatogram. The doubled peaks that begin four bases 5' to the start of the gRNA sequence indicate that this founder mouse was genetically mosaic for two distinct deletions between the exon 1 and exon 25 gRNA sequences. The two distinct deletions were resolved using Poly Peak Parser. (**C**) Mutant *Mtmr5*$^{-/-}$ mouse #992 (F1 progeny of mouse #684) sequence with coding frame indicated and premature stop codon identified (*).

**Supplemental Figure S2:** Mtmr5 protein levels from wild type, *Mtmr5*$^{+/-}$, and *Mtmr5*$^{-/-}$ mice, analyzed by immunoblotting. The protein loading controls were Memcode total protein stain and GAPDH. (**A**) At P7, a protein band with an apparent molecular mass of 75-100 kDa was detected in *Mtmr5*$^{-/-}$ mouse brains using a C-terminal anti-Mtmr5 antibody. Both short and long exposure of Mtmr5 immunoblot images are shown. The truncated Mtmr5 (predicted to be 84.8 kDa) had a protein level of 0.17 a.u. relative to the level of the full-length protein, which had a mean level of 1.03 ± 0.04 a.u. (n = 2 wild type, and n = 1 *Mtmr5*$^{-/-}$; mean ± SD). (**B**) At 5 months, the truncated Mtmr5 protein was not detected in mutant mouse brains (n = 3 wild type, n = 1 *Mtmr5*$^{+/-}$, and n = 3 *Mtmr5*$^{-/-}$ mice; *n.s.*, non-specific protein bands). (**C**) An increased exposure of the Mtmr5 immunoblot from Fig. 2C of sciatic nerve lysates from 5-month-old mice. The ~85 kDa truncated Mtmr5 protein was not detected (n = 3 wild type, n = 3 *Mtmr5*$^{+/-}$, and n = 3 *Mtmr5*$^{-/-}$ mice; *n.s.*, non-specific protein bands).

**Supplemental Figure S3:** Reduced weight in the absence of Mtmr5. (**A & B**) *Mtmr5*$^{-/-}$ male and female mice alongside sex-matched littermate controls at postnatal day 38 (P38). (**C**) At P38, *Mtmr5*$^{-/-}$ male mice were significantly smaller than controls. Average weights: 20.33 ± 1.03 g, 20.02 ± 2.57 g, and 14.47 ± 2.17 g, for *Mtmr5*$^{+/+}$, *Mtmr5*$^{+/-}$, and *Mtmr5*$^{-/-}$, respectively (n = 8, 13, and 6, respectively). (**D**) Female *Mtmr5*$^{-/-}$ mice were significantly smaller than controls at P38. Average weights: 16.94 ± 1.69 g, 17.19 ± 1.30 g, and 12.15 ± 0.98 g for *Mtmr5*$^{+/+}$, *Mtmr5*$^{+/-}$, and *Mtmr5*$^{-/-}$, respectively (n = 7, 14, and 8, respectively) (mean ± SD; one-way ANOVA with post-hoc Tukey test; ***$p \leq 0.001$; ****$p \leq 0.0001$; *n.s.*, not significant).

**Supplemental Figure S4:** No apparent defects in axon guidance or nerve branching in *Mtmr5*$^{-/-}$; *Mtmr13*$^{-/-}$ embryos at E13. (**A**) Representative images of *Mtmr5*$^{+/+}$; *Mtmr13*$^{-/-}$ (control) and *Mtmr5*$^{-/-}$; *Mtmr13*$^{-/-}$ (dKO) embryos that had been immuno-labeled with neurofilament medium chain (NF-M) antibody to reveal axons. Scale bar: 2000 µm. (**B-C**) Corresponding higher magnification images of *Mtmr5*$^{+/+}$; *Mtmr13*$^{-/-}$ (control) and dKO embryo ears. Two major nerve branches, the great auricular nerve and the auriculotemporal nerve, which develop between E12 and E13 in the mouse pinna, are indicated (*red circles*). Defects in axon guidance and nerve branching, including those caused by genetic mutations, can be discerned in these nerves by E13 (1). dKO animals showed normal nerve branching patterns and did not show evidence of abnormal axon guidance. (**D-F**) Additional examples of littermate control (*Mtmr5*$^{+/-}$; *Mtmr13*$^{-/-}$) and dKO (*Mtmr5*$^{-/-}$; *Mtmr13*$^{-/-}$) embryos are shown. Scale bar: 200 µm.
